# Targeting AGTR1/NF-κB /CXCR4 axis by miR-155 attenuates oncogenesis in Glioblastoma

**DOI:** 10.1101/815068

**Authors:** Anukriti Singh, Nidhi Srivastava, Bushra Ateeq

## Abstract

Glioblastoma (GBM) represents the most aggressive malignancy of the brain. Angiotensin II Receptor Type 1 (AGTR1) upregulation has been associated with proliferative and infiltrative properties of glioma cells. However, the underlying mechanism of AGTR1 upregulation in GBM is still unexplored. To understand the post-transcriptional regulation of *AGTR1* in GBM, we screened 3’untranslated region (3’UTR) of *AGTR1* by using prediction algorithms for binding of miRNA. Interestingly, miR-155 showed conserved binding on the 3’UTR of *AGTR1*, subsequently confirmed by *AGTR1*-3’UTR-luciferase reporter assay. Furthermore, stable miR-155 overexpressing GBM cells show decrease in AGTR1-mediated cell proliferation, invasion, foci formation and anchorage-independent growth. Strikingly, immunodeficient mice implanted with stable miR-155 overexpressing SNB19 cells show remarkable reduction (∼95%) in tumor burden compared to control. Notably, miR-155 attenuates NF-κB signaling downstream of AGTR1 leading to reduced CXCR4 and AGTR1 levels. Mechanistically, miR-155 mitigates AGTR1-mediated, angiogenesis, epithelial-to-mesenchymal transition, stemness, ERK/MAPK signaling and promotes apoptosis. Similar effects in cell-based assays were observed by using pharmacological inhibitor of IκB Kinase (IKK) complex. Taken together, we established that miRNA-155 post-transcriptionally regulates *AGTR1* expression, abrogates AGTR1/NF-κB/CXCR4 signaling axis and elicits pleiotropic anticancer effects. This study opens new avenues for using IKK inhibitors and miRNA-155 replacement therapies for the treatment of AGTR1-positive malignancies.

## Introduction

Glioblastoma (GBM) is the most common leading cause of cancer death among all adult primary brain tumors. Regardless of the recent developments in GBM treatment and therapeutics comprising maximum safe resection, followed by radio-chemotherapy, the average patient survival time is less than 15 months [1, 2]. Surgical treatments present many limitations as the tumor infiltrates and diffuses in the surrounding parenchyma leading to disease relapse [3]. GBM has been categorized into proneural, neural, classical and mesenchymal subtypes; the mesenchymal subgroup has been linked to the worst prognosis [4]. Genetic alterations such as MGMT promoter methylation and *IDH1* mutation are known to be associated with therapeutic resistance and recurrence [5]. The intratumoral heterogeneity of GBM poses several challenges such as post-therapy resistance and compromised responses to radio-chemotherapy [6]. Therefore, there is a need to investigate the molecular mechanisms underlying poor prognosis of GBM, and develop novel therapeutic approaches based on targeting specific molecular subtypes.

Increased levels of Angiotensin II Receptor Type 1 (AGTR1), a G-protein coupled receptor has been linked to high grade astrocytoma and poor patient prognosis [7]. Moreover, GBM cell lines overexpressing AGTR1 and Angiotensin II Receptor Type 2 (AGTR2) exhibit a mitogenic response upon treatment with angiotensin peptides [8]. Furthermore, AGTR1 overexpression has also been linked to multiple human malignancies including ovarian, renal, pancreatic, breast, and brain [9–12]. While in breast cancer (BCa), significant overexpression of AGTR1 represents a subpopulation of Human epidermal growth factor (HER2)-negative and estrogen receptor (ER) positive patients [13], in ovarian cancer its high expression has been linked to metastasis by promoting multicellular spheroid formation [14]. Mechanistically, overexpression of AGTR1 has been observed to inhibit apoptosis while promoting angiogenesis, cell proliferation and inflammation [15, 16]. Conversely, blockage of AGTR1 in C6 rat glioma cells results in attenuated angiogenesis, cell proliferation and tumor progression. [12]. Although, AGTR1 has been deemed as an oncogene in various cancers, its role in GBM is yet to be explored completely. Further, the underlying mechanism of its overexpression in GBM is not well understood and needs to be elucidated.

One of the mechanisms for AGTR1-mediated tumorigenesis has been attributed to multiple signaling pathways downstream of AGTR1, for instance, nuclear factor-κB (NF-κB) signaling in BCa [17]. Moreover, angiotensin II (ATII), the ligand of AGTR1 is known to activate NF-κB signaling downstream of AGTR1 in endothelial and vascular smooth muscle cells [18]. The IκB Kinase (IKK) complex is important for the positive regulation of NF-κB, which by phosphorylation-induced, proteasome-mediated degradation of IκB results in nuclear accumulation of NF-κB, wherein it binds to its target genes involved in cell survival, growth, immune response and inflammation [19]. Activation of NF-κB signaling in GBM often triggers multiple downstream targets, such as activation of GPCR named C-X-C chemokine receptor type 4 (CXCR4) [20], which promotes cell migration and metastases [21, 22]. CXCR4 is a major chemokine receptor, and a known marker for GBM metastasis. It mediates the cell survival, invasion [23, 24] and proliferation of GBM progenitor cells [25, 26], deeming it as a promising therapeutic target in GBM.

In this study, we discovered the mechanism underlying AGTR1 overexpression in GBM and its downstream signaling pathways associated with AGTR1-mediated oncogenesis. Further, we established the role of miR-155 in the post-transcriptional regulation of *AGTR1* in GBM. We also showed that activation of AGTR1 via ATII stimulation in GBM cells activate NF-κB signaling leading to CXCR4 overexpression as well as AGTR1 upregulation, thereby forming a positive feedback regulatory loop. Moreover, ectopic expression of miR-155 in GBM cells attenuates AGTR1 downstream signaling thereby disrupting this regulatory loop. Alternatively, targeting NF-κB signaling by an IKK complex inhibitor, results in downregulation of AGTR1 and CXCR4 expression, leading to reduced AGTR1-mediated oncogenicity. Conclusively, this study reveals a novel regulatory mechanism involving miR-155, which targets AGTR1/NF-κB/CXCR4 axis and abrogates GBM progression.

## Results

### Silencing AGTR1 decreases oncogenesis in glioblastoma

To examine the expression of *AGTR1* in GBM, we analyzed TCGA-GBM (The Cancer Genome Atlas-Glioblastoma) and Sun Brain (GSE4290) patients’ cohorts. We observed that both cohorts exhibit increased expression of *AGTR1* in GBM tumors with respect to normal tissue (Fig. 1a, b). We next evaluated the overall survival probability of GBM patients with high *versus* low *AGTR1* expression. Interestingly, patients with high *AGTR1* expression show overall low survival probability compared to the patients with low *AGTR1* levels (Fig. 1c), implicating an association between elevated AGTR1 levels with poor survival of the clinically advanced GBM patients. Several independent studies implicated AGTR1 upregulation in cell proliferation, invasion and distant metastases in multiple malignancies [7, 13, 17]. Therefore, to ascertain the role of AGTR1 in GBM oncogenesis, we performed stable shRNA-mediated knockdown of *AGTR1* in SNB19 cells, an AGTR1-positive GBM cell line (Fig. 1d, e), and characterized them for oncogenic properties. Importantly, a significant decrease in the cell proliferation of SNB19-shAGTR1 cells was observed with respect to control (Fig. 1f). Similarly, a marked decrease in the migratory and invasive potential was observed in SNB19-shAGTR1 cells (∼ 80% and 85% respectively) as compared to control (Fig. 1g, h). Moreover, a significant reduction (∼60%) in the foci forming ability of these cells was also noted (Fig. 1i). These results confirm the oncogenic role of AGTR1 in GBM as silencing of *AGTR1* successfully abrogated oncogenic properties of GBM cells

**Fig. 1.**
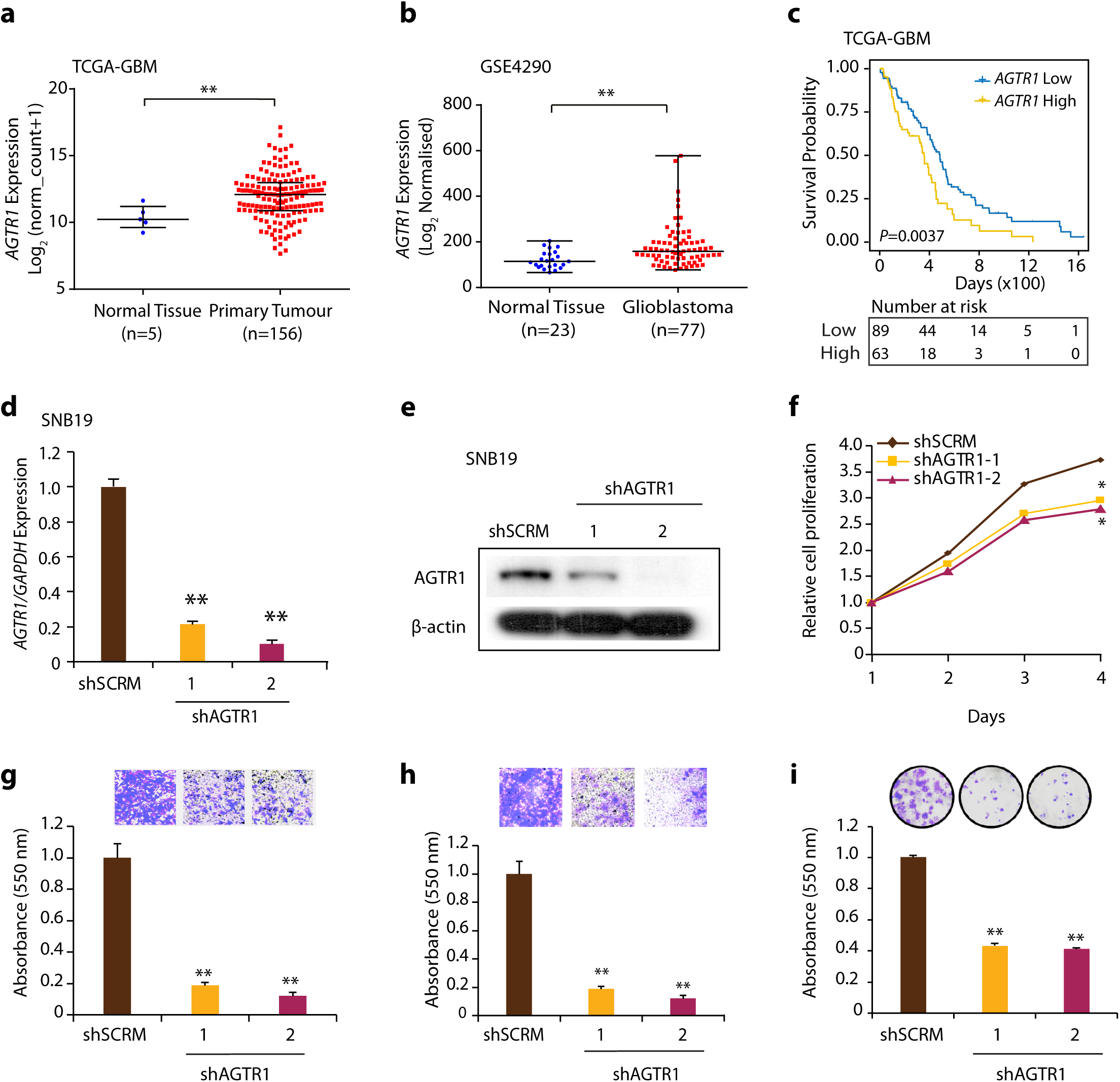
Silencing *AGTR1* leads to reduced oncogenesis in glioblastoma. **a** Scatter plot showing expression of *AGTR1* in normal tissue (n=5) and primary tumors (n=156) in the TCGA-GBM cohort. **b** Scatter plot showing expression of *AGTR1* in normal tissue (n=23) and glioblastoma patients (n=77) in the GSE4290 cohort. **d** Bar plot showing expression of *AGTR1* in SNB19-shAGTR1 and SNB19-shSCRM cells by quantitative PCR. **e** Immunoblot assay for AGTR1 levels in the same cells as **d**. β-actin was used as a loading control. **f** Line graph showing cell proliferation assay using *AGTR1*-silenced (shAGTR1) and control (shSCRM) SNB19 cells. **g** Bar plot depicting Boyden chamber migration assay using the same cells as in **d**. Inset displaying representative fields of migrated cells (n=3 biologically independent samples; data represent mean ± SEM). **h** Bar plot depicting Boyden chamber Invasion assay using the same cells as in **d**. Inset displaying representative fields of invaded cells (n=3 biologically independent samples; data represent mean ± SEM). **i** Bar plot showing foci formation assay using the same cells as in **d**. Inset showing representative images of foci (n=3 biologically independent samples; data represent mean ± SEM). Data represent mean ± SEM. \**P ≤* 0.05, ** *P ≤* 0.001 using two-tailed unpaired Student’s *t-*test.

### Ectopic expression of miR-155 leads to decreased AGTR1-mediated oncogenesis

We assessed the role of miRNAs in the post-transcriptional regulation of AGTR1, by employing miRNA prediction algorithms, namely TargetScan [27], miRanda [28] (microRNA.org), MicroT4 [29] (DIANA tools), RNA22 [30] as well as experimentally supported database TarBase [31], and screened for putative miRNAs binding sites on the 3’-UTR of *AGTR1* transcript. Interestingly, miR-155 was independently predicted to have a consensus binding site on the 3’UTR of *AGTR1* by all the five algorithms (Fig. 2a, b). Next, to validate the miR-155 binding to the 3’-UTR of *AGTR1*, we generated 3’-UTR wild type (3’-UTR-WT) and mutant (3’-UTR-mut) *AGTR1 Firefly/Renilla* dual-luciferase reporter constructs, and co-transfected them with synthetic mimic for miR-155 in HEK293T cells. A significant decrease in luciferase reporter activity was recorded in 3’-UTR-WT transfected HEK293T cells, while no reduction in luciferase activity was observed with 3’-UTR-mut construct (Fig. 2c). We next evaluated the expression of miR-155 and *AGTR1* in GBM cell lines (SNB19 and U138), and an inverse correlation between the expression patterns of *AGTR1* and miR-155 was observed in both cell lines (Fig. 2d). Further, to ascertain the miR-155-mediated negative regulation of *AGTR1*, we established stable miR-155 overexpressing SNB19 cells and analyzed the expression of *AGTR1*. Remarkably, a significant reduction in the AGTR1 levels was observed both at transcript and protein levels in SNB19-miR-155 overexpressing cells (Fig. 2e). Our findings demonstrate that miR-155 binds to 3’-UTR of *AGTR1* and leads to its post-transcriptional repression. We next investigated the functional significance of miR-155 using stable miR-155 overexpressing SNB19 cells (SNB19-miR-155) and examined for any alteration in their oncogenic properties. Notably, a significant decrease in cell proliferation was observed in SNB19-miR-155 cells compared to control (Fig. 2f). Similarly, a significant reduction in cell migration (∼ 80%) and invasion (∼ 60%) was observed in SNB19-miR-155 cells (Fig. 2g, h). Notably, this change in migration and invasion of SNB19-miR-155 cells was not due to a decrease in the proliferation rate of these cells, as invasion and migration assays were terminated at 24 and 16 hours, respectively. Furthermore, to analyze the oncogenic transformation of SNB19-miR-155 cells, foci formation assay was performed, and a significant reduction in the number of foci was observed (Fig. 2i). Moreover, a striking decrease in anchorage-independent growth was recorded in SNB19-miR-155 overexpressing cells (∼ 50-80%), whereas a moderate reduction in the number of soft agar colonies was observed in SNB19-miR-155 pool cells (∼15%) in comparison to control cells, while remarkable reduction in the size of the soft agar colonies was evident in both pool as well as single clones (Fig. 2j). To further examine the effect of miR-155 on tumorigenesis, we subcutaneously implanted SNB19-miR-155 and SNB19-CTL cells in the flank region of immunodeficient NOD/SCID mice, and tumor growth was monitored. Intriguingly, a remarkable regression (∼95%) in tumor burden of mice bearing SNB19-miR-155 overexpressing xenografts was observed, compared to control the group (Fig. 2k). Conclusively, our findings implicate the tumor-suppressive role of miR-155 in GBM by downregulating *AGTR1* expression and abrogating AGTR1-mediated tumorigenesis.

**Fig. 2.**
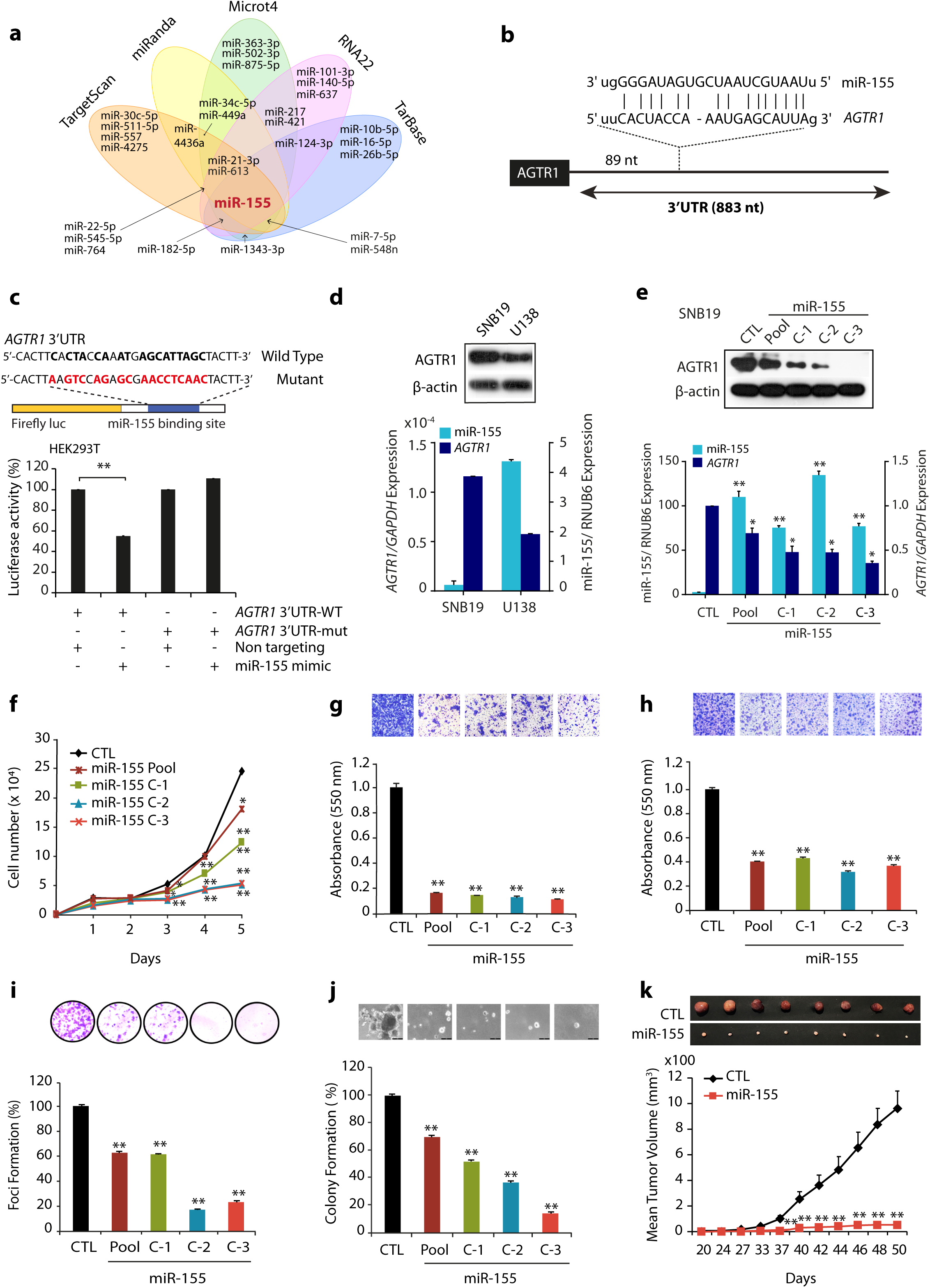
miR-155 post-transcriptionally regulates *AGTR1* and its overexpression abrogates oncogenic properties of glioblastoma cells. **a** Venn diagram displaying computationally predicted miRNAs targeting *AGTR1* by TargetScan, miRanda, Microt4, RNA22, and TarBase. **b** Schema displaying the predicted binding site of miR-155 on the 3’UTR of *AGTR1*. **c** Schema showing luciferase reporter construct with wild-type or mutated (red residues with alteration) *AGTR1* untranslated region (3’UTR) downstream of firefly luciferase reporter gene (top panel). Bar plot showing luciferase reporter activity in HEK293T cells co-transfected with miR-155 mimics and mutant or wild-type 3’UTR *AGTR1* constructs. **d** Quantitative PCR data showing expression of *AGTR1* and miR-155 in glioblastoma cell lines. The bar plot depicts *AGTR1* and miR-155 as 2^-dCt. Immunoblot assay showing AGTR1 levels in glioblastoma cell lines. β-actin was used as a loading control (top panel). **e** Same as **d**, except SNB19-miR155 and SNB19-CTL cells were used. Top panel showing immunoblot for AGTR1 levels using the same cells. β-actin was used as a loading control. Data represent mean ± SEM. \**P ≤* 0.05, ** *P ≤* 0.001 using two-tailed unpaired Student’s *t-*test. **f** Line graph showing cell proliferation assay using SNB19-miR155 cells and SNB19-CTL cells. **g** Bar plot depicting Boyden chamber migration assay using the same cells as in **f**. Inset displaying representative fields of migrated cells (n=3 biologically independent samples; data represent mean ± SEM). **h** Bar plot depicting Boyden chamber invasion assay using the same cells as in **f**. Inset displaying representative fields of invaded cells (n=3 biologically independent samples; data represent mean ± SEM). **i** Foci formation assay using the same cells as in **f**. Inset showing representative images of foci (n=3 biologically independent samples; data represent mean ± SEM). **j** Soft agar colony formation assay using the same cells as in **f**. Inset displaying representative soft agar colonies (n=3 biologically independent samples; data represent mean ± SEM). **k** Line graph showing mean tumor growth in NOD/SCID mice (n=8 per group) subcutaneously implanted with SNB19-CTL and SNB19-miR-155 cells. Data represent mean ± SEM. \**P ≤* 0.05, ** *P ≤* 0.001 using two-tailed unpaired Student’s *t-*test.

### Global gene expression profile of miR-155 reveal multiple deregulated oncogenic biological processes

To investigate the biological pathways leading to the decreased oncogenic potential of SNB19-miR-155 cells, we performed global gene expression profiling of SNB19-miR-155 and SNB19-CTL cells. Analysis of gene expression profiles of these cells show 735 genes to be downregulated (log_2_ fold change ≤ -0.6 and *P ≤* 0.05) and 638 genes to be upregulated (log_2_ fold change ≤ 0.6 and *P≤* 0.05) in SNB19-miR-155 cells with respect to control cells (Supplementary Table S1). We next employed DAVID (Database for Annotation, Visualization and Integrated Discovery), GSEA (Gene Set Enrichment Analysis), and Enrichment Map analysis (Supplementary Fig. S1) to examine the biological processes that were differentially regulated upon ectopic expression of miR-155 in GBM cells. Interestingly, majority of genes downregulated upon miR-155 overexpression were associated with CXCR4 signaling pathway, positive regulation of MAPK, ERK1/2 cascade, blood vessel development and positive regulation of cell migration. While, genes associated with programmed cell death, apoptotic processes, negative regulation of cell proliferation and negative regulation of vasculature development were upregulated upon miR-155 overexpression (Fig. 3a). Additionally, GSEA of miR-155 overexpressing cells showed a significant reduction in the enrichment of genes associated to the classical subtype of GBM, cell-cycle checkpoint, and stemness (Fig. 3b). Furthermore, our GSEA output shows a significant reduction in the expression of genes related to epithelial-to-mesenchymal transition (EMT) (Fig. 3b). Diminished expression of genes associated with EMT in GBM was also observed in our gene expression data (Fig. 3c) We next validated this data by examining the expression of genes involved in EMT, and a significant increase in E-Cadherin (*CDH1*), an epithelial marker was observed with a concomitant decrease in mesenchymal markers namely Vimentin (*VIM)* and SLUG (*SNAI2*) (Fig. 3c). Similarly, a significant increase in the E-Cadherin and a decrease in Vimentin and N-cadherin at the protein level was noted (Fig. 3d). Since, both MAPK and Akt signaling pathways play a key role in regulating cell proliferation and survival, we therefore observed the phosphorylation status of ERK (p-ERK) and Akt (p-Akt) as a readout of these pathways. As apparent in the DAVID analysis, a significant reduction of the p-ERK and p-Akt levels in SNB19-miR-155 cells was detected with respect to control (Fig. 3e). Furthermore, apoptosis was another key pathway delineated by our gene expression analysis, hence we explored the expression of Bcl-2 and Bcl-XL proteins, known to inhibit cell death [27]. Intriguingly, we found a remarkable decrease in Bcl-2 and Bcl-XL (anti-apoptotic proteins) in SNB19-miR-155 cells compared to control cells (Fig. 3f). Aldehyde dehydrogenase (ALDH) is a known stem cell marker for GBM [28], we therefore performed ALDH assay, and a significant decrease (∼67%) in its activity in SNB19-miR-155 cells compared to control was recorded (Fig. 3g). Further, we analyzed the expression of *KIT*, targeting which is known to reduce GBM cell viability in vitro [29]. We also examined the expression of important treatment resistant tumor initiating cells’ and glioma stem cells’ markers *CD24* and CD44. [30]. We observed a significant decrease in expression of *CD24* (70-30%) and *KIT* (45-30%) (Fig. 3h) and percentage CD44 positive cells (∼60%) in miR-155 overexpressing SNB19 cells compared to control (Fig. 3i). As prominent in our GSEA analysis, we also carried out cell cycle analysis of SNB19 cells transiently transfected with miRNA mimics and observed a marked increase in S phase cell arrest (Fig. 3j). Conclusively, our findings show that miR-155 downregulates the expression of genes associated with various oncogenic pathways namely, survival, ERK/MAPK signaling, EMT, stemness, promotes apoptosis and induces cell cycle arrest, signifying that miR-155 replacement strategies may offer an alternative therapeutic intervention for the glioblastoma patients.

**Fig. 3.**
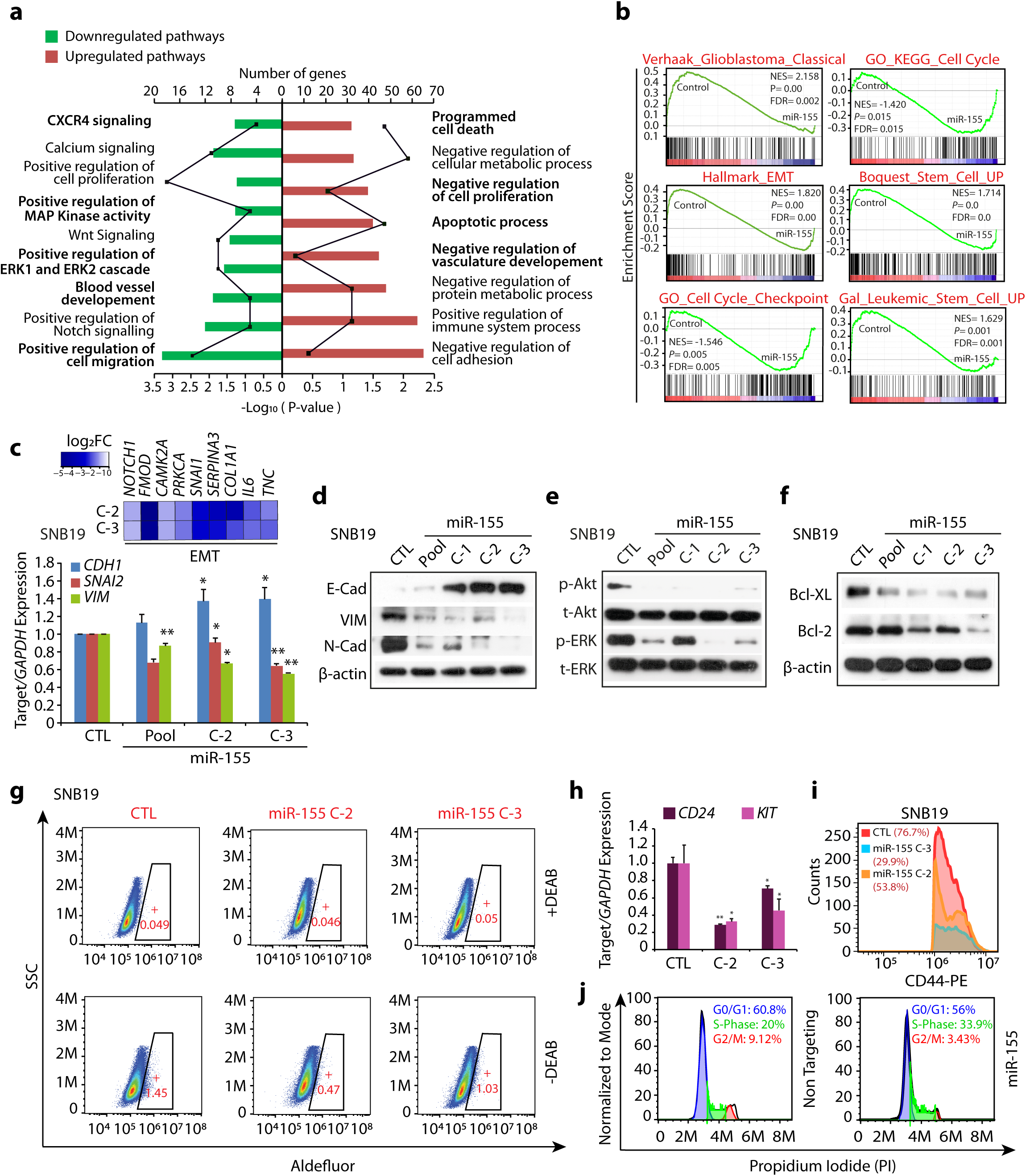
MiR-155 abrogates multiple oncogenic pathways in glioblastoma. **a** Bar plot showing DAVID analysis of gene expression profile data of SNB19-miR-155 cells compared to SNB19-CTL cells, showing downregulated (green) and upregulated (red) pathways. Frequency polygon (line in black) represents the number of genes and the bars represent –Log_10_ (*P* values). **b** The Gene Set Enrichment Analysis (GSEA) plots depicting deregulated processes using same gene expression profile data as in **a**. **c** Heatmap depicting the relative expression of genes related to EMT in the same cells as **a** (top panel). Shades of blue represent log_2_ fold change in gene expression. Bar plot showing quantitative PCR data for the relative expression of EMT markers in the same cells as **a**. **d** Immunoblot assays for E-Cadherin, Vimentin, and N-Cadherin using the same cells as in **a**. β-actin was used as a loading control. **e** Immunoblot assays for phospho (p), total (t) Akt and ERK1/2 using the same cells as in **a**. **f** Immunoblot assays for anti-apoptotic proteins Bcl-2 and Bcl-XL for the same cells as in **a**. **g** Aldeflour assay showing aldehyde dehydrogenases (ALDH) expression using SNB19-miR-155 (C-2 and C-3) and SNB19-CTL cells. The graphs depict the fluorescence intensity of the catalysed ALDH substrate in the presence and absence of DEAB (ALDH activity inhibitor). Windows marked in graphs represent the percentage of ALDH positive cell population. **h** Bar plot showing quantitative PCR data for the relative expression of *CD24* and *KIT* in the same cells as **g**. **i** Flow cytometry analysis showing CD44 expression using the same cells as **g**. **j** Flow cytometry analysis depicting S-phase cell cycle arrest of SNB19 cells transfected with miR-155 mimics stained with propidium iodide (PI). Data represent mean ± SEM. \**P ≤* 0.05, ** *P ≤* 0.001 using two-tailed unpaired Student’s *t-*test.

### MiR-155 mediates its tumor-suppressive effects by downregulating angiogenesis

Angiogenesis or formation of new blood vessels from the pre-existing vasculature, is crucial for the growth and survival of all cancers including GBM. Furthermore, an increase in progression-free survival was observed in GBM patients administered with bevacizumab, an angiogenesis inhibitor, in phase III clinical trials [31, 32]. Our Enrichment Map as well as downregulated pathway analysis show a decrease in the expression of genes involved in angiogenesis and Vascular endothelial growth factor (VEGF) signaling in miR-155 overexpressing cells (Supplementary Fig. S1and Fig. 4a, b). Therefore, we sought to investigate the effect of miR-155 on angiogenesis and examined the expression of genes associated with angiogenesis in GBM. Interestingly, a significant reduction in the expression of established angiogenesis markers, namely, Nestin (*NES*) [33], Atypical chemokine receptor 3 (*ACKR3*) [34], Platelet derived growth factor β (*PDGFRB*) [35], and Anterior grade protein 2 (*AGR2*) [36] was observed (Fig. 4c). To investigate the angiogenic potential of miR-155 in GBM, we implanted SNB19-miR-155 or control cells in chick chorioallantoic membrane (CAM) assay, a popular *in vivo* model for studying angiogenesis. Intriguingly, our CAM assay data indicates a remarkable reduction in angiogenesis in the CAMs implanted with SNB19-miR155 cells as compared to the control (Fig 4d). We also measured the features of angiogenic vessels by using ImageJ angiogenesis analyzer [37], and a significant reduction in the number of nodes, branches, meshes, junctions, segments along with mean mesh size, total segments length and total branches length was observed in SNB19-miR-155 implanted CAM group compared to control (Fig. 4e). Taken together, we conclude that overexpression of miR-155 attenuates angiogenesis in AGTR1-positive SNB19 GBM cells, thus establishing the importance of miR-155 as a therapeutic intervention against tumor angiogenesis for glioblastoma.

**Fig. 4.**
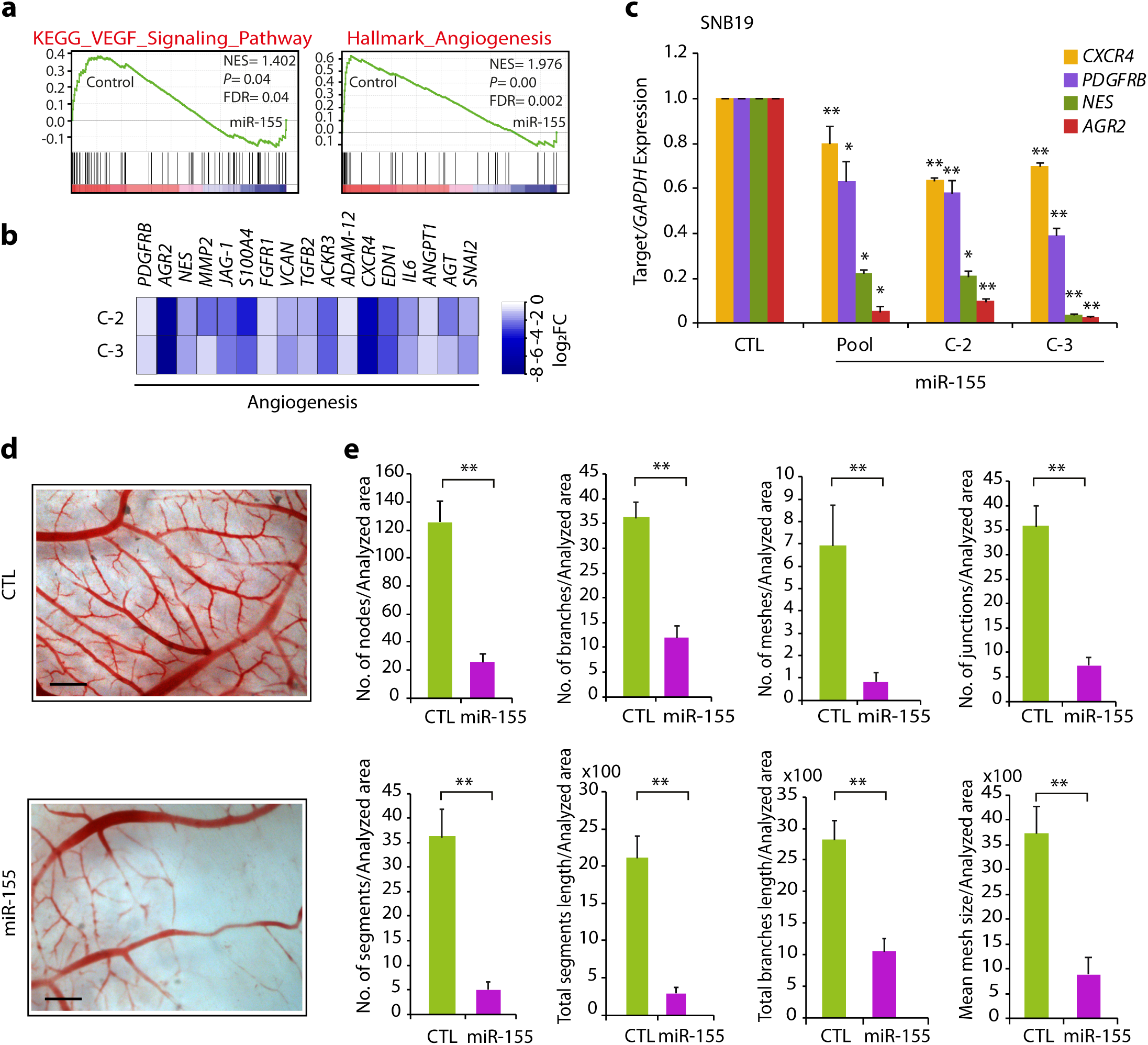
MiR-155 attenuates neovascularization in chorioallantoic membrane (CAM) assay. **a** The Gene Set Enrichment Analysis (GSEA) plots depicting angiogenesis and VEGF signaling to be enriched and their corresponding statistical metrics in SNB19-miR155 and control cells. **b** Heatmap depicting the relative expression of genes related to angiogenesis using the same cells as in **a.** Shades of blue represent log_2_ fold change in gene expression. **c** Bar plots showing quantitative PCR data for the relative expression of angiogenesis markers using the same cells as in **a**. **d** Representative images of chick CAM assay using the same cells as in **a**. Scale bar represents 1cm. **e** Bar plots representing angiogenic properties of the CAM assay using microscopic images analyzed by Angiogenesis analyzer in Image J. Data represent mean ± SEM. \**P ≤* 0.05, ** *P ≤* 0.001 using two-tailed unpaired Student’s *t-*test.

### Disrupting the AGTR1/NF-κB/CXCR4 axis attenuates oncogenesis in Glioblastoma

The expression of C-X-C chemokine receptor type 4 (CXCR4) is known to mediate tumor invasion and targeting this receptor has been shown to reduce the invasive properties of GBM cells, sensitizing them towards radiation-induced apoptosis [38]. Importantly, the CXCR4 signaling pathway emerged as one of the most significant biological processes downregulated upon miR-155 overexpression in GBM by DAVID and GSEA analysis (Fig. 3a and Fig. 5a). Hence, we next analyzed TCGA-GBM and Sun Brain (GSE4290) cohorts for the expression of *CXCR4* and found it to be significantly overexpressed in both GBM patients’ cohorts (Supplementary Fig. S2a, b). Moreover, TCGA-GBM patients with higher expression of *CXCR4* show reduced overall survival probability compared to patients with low *CXCR4* expression (Supplementary Fig. S2c), suggesting that its higher expression is associated with poor patient survival. Hence, we next examined the expression of CXCR4 in SNB19-miR-155 cells, and a concomitant decrease in the CXCR4 expression was observed with respect to control (Fig. 5b). To pinpoint the exact mechanism involved in reduced CXCR4 expression, we employed miRNA prediction tools to examine whether miR-155 binds on the 3’UTR of the *CXCR4*; nevertheless, absence of miR-155 binding indicates that some alternate regulatory mechanism might be involved. Interestingly, our GSEA data revealed both CXCR4 and NF-κB signaling pathways to be significantly downregulated in SNB19-miR155 cells compared to control (Fig. 5a). NF-κB signaling being a master regulator of cell survival, immunity, inflammation, as well as considering its critical role in the maintenance of cancer stem-like cells and resistance to radiotherapy in GBM [20], we sought to investigate the possible link between miR-155 and NF-κB signaling. Previously, NF-κB signaling is known to induce CXCR4 expression in BCa cells [21] and upon hepatocyte growth factor stimulation in glioma [39], hence we conjectured that the decrease in CXCR4 expression in miR-155 overexpressing cells might be influenced by NF-κB signaling. Further, it has been shown that ATII activates NF-κB and CREB signaling via p38 MAPK leading to the upregulation of AGTR1 [40], hence we speculated that NF-κB signaling might be involved in upregulation of AGTR1 in GBM. To examine this, we looked at the phosphorylation of p65 subunit (p-p65) levels upon ATII stimulation, a bonafide ligand for AGTR1. As expected, higher p-p65 levels with a concomitant increase in AGTR1 and CXCR4 expression was observed in ATII stimulated SNB19 cells (Supplementary Fig. S2d, e, and Fig. 5c). Conversely, stimulating SNB19-miR-155 cells with ATII resulted in a robust decrease in the p-p65 levels compared to SNB19-CTL cells (Fig. 5d). Similarly, a marked decrease in AGTR1 and CXCR4 levels was also noted in SNB19-miR-155 cells (Fig. 5d). Our data suggest that ATII stimulation activates NF-κB signaling, which in turn results in increased expression of CXCR4 and AGTR1 in GBM cells, while the presence of miR-155 disrupts this feedback regulatory loop.

**Fig. 5.**
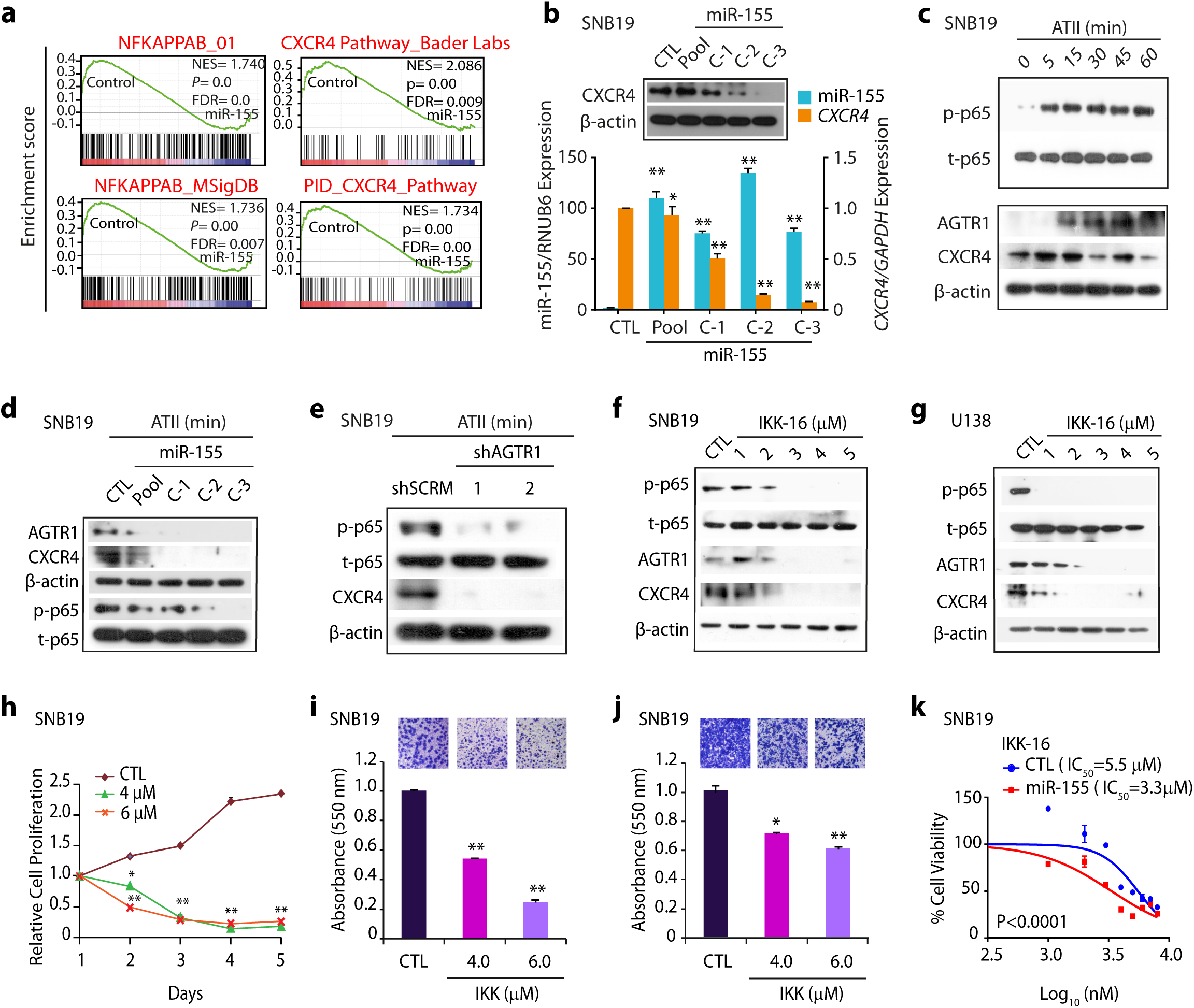
Targeting AGTR1 in Glioblastoma results in reduced NF-κB signaling and CXCR4 expression. **a** Gene Set Enrichment Analysis (GSEA) for NF-κB and CXCR4 pathway and their associated statistical metrics in SNB19-miR-155 and SNB19-CTL cells. **b** Bar plot showing relative expression of CXCR4 and miR-155 by quantitative PCR using the same cells as in **a**. Immunoblot assay for CXCR4 levels using the same cells. β-actin was used as a loading control. **c** Immunoblot showing phospho (p) and total (t) p65, AGTR1 and CXCR4 levels using ATII stimulated SNB19 cells at indicated time points. β-actin was used as a loading control. **d** Immunoblot assays for AGTR1, CXCR4, phospho (p) and total (t) p65 levels using SNB19-miR-155 and SNB19-CTL cells stimulated with ATII for 15 minutes. β-actin was used as a loading control. **e** Immunoblots for phospho (p) and total (t) p65 and CXCR4 levels using SNB19-shAGTR1 and SNB19-shSCRM cells stimulated with ATII for 15 minutes. β-actin was used as a loading control. **f** Immunoblots for phospho (p) p65, total (t) p65, AGTR1 and CXCR4 levels using ATII stimulated (15 minutes) and IKK-16 treated SNB19 cells. β-actin was used as a loading control. **g** Same as in **f** except U138 cells were used. β-actin was used as a loading control. **h** Line graph showing cell proliferation assay for IKK-16 treated SNB-19 cells at the indicated concentration. **i** Bar plot depicting Boyden chamber invasion assay using the same cells as in **i**. Inset showing representative fields of invaded cells (n=3 biologically independent samples; data represent mean ± SEM). **j** Bar plot depicting Boyden chamber migration assay using the same cells as in **i**. Inset showing representative fields of migrated cells (n=3 biologically independent samples; data represent mean ± SEM). **k** Dose-response curve with a range of IKK-16 using SNB19-miR-155 and SNB19-CTL cells. IC_50_ values were calculated using GraphPad Prism software. Data represent mean ± SEM. \**P ≤* 0.05, ** *P ≤* 0.001 using two-tailed unpaired Student’s *t-*test.

To further corroborate our finding that miR-155 disrupts AGTR1/NF-κB /CXCR4 axis in GBM, we stimulated SNB19-shAGTR1 cells with ATII and examined for NF-κB signaling. Interestingly, ATII stimulation in SNB19-shAGTR1 cells shows decreased p-p65 levels accompanied with a notable reduction in the CXCR4 expression compared to control SNB19-shSCRM cells (Fig. 5e). Collectively, silencing *AGTR1* in GBM cells demonstrates similar results as achieved via miR-155 overexpression, thereby providing compelling evidence that the downregulation of AGTR1 results in reduced NF-κB signaling. Finally, our data indicate that targeting AGTR1 either by miR-155 or RNA-interference could abrogate NF-κB downstream signaling in AGTR1-positive GBM subtype.

Next, we sought to examine whether abrogating NF-κB signaling by specific pharmacological inhibitor, could lead to reduction in AGTR1 and CXCR4 expression and associated oncogenic properties. We used IKK-16, a selective IκB kinase (IKK) inhibitor for, IKK-1, IKK-2 and IKK complex [41]. Interestingly, a robust decrease in p-p65 levels was observed in ATII stimulated SNB19 cells treated with IKK-16 (Fig. 5f). Moreover, a marked decrease in both AGTR1 and CXCR4 expression was also noted in these cells (Fig. 5f). Likewise, a remarkable decrease in p-p65 levels, AGTR1, and CXCR4 expression was observed in U138 GBM cells (Fig. 5g).

Next, we also investigated any alteration in oncogenic properties by abrogating the NF-κB signaling pathway via IKK-16, a significant decrease in cell proliferation was observed in ATII stimulated and IKK-16 treated SNB19 cells compared to vehicle control (Fig. 5h). Similarly, a marked reduction in invasion and migration was recorded in IKK-16 treated SNB19 cells post ATII stimulation (Fig. 5i, j). Furthermore, we assessed change in drug-sensitivity in miR-155 overexpressing GBM cells and found that SNB19-miR-155 show enhanced chemosensitivity towards IKK-16 compared to SNB19-CTL cells (Fig. 5k), which leads us to infer that overexpressing miR-155 may be useful to overcome drug resistance in GBM. Conclusively, our findings implicate the role of miR-155 in abrogating NF-κB signaling downstream of AGTR1, thereby leading to inhibition of CXCR4 expression and AGTR1 feedback upregulation (Fig. 6). Thus, nominating miR-155 as a key player in therapy for AGTR1-positive GBM. Hence, disrupting AGTR1/NF-κB/CXCR4 axis by miR-155 or using pharmacological inhibitors against NF-κB signaling might be beneficial for treating AGTR1-positive subset of GBM patients.

**Fig. 6.**
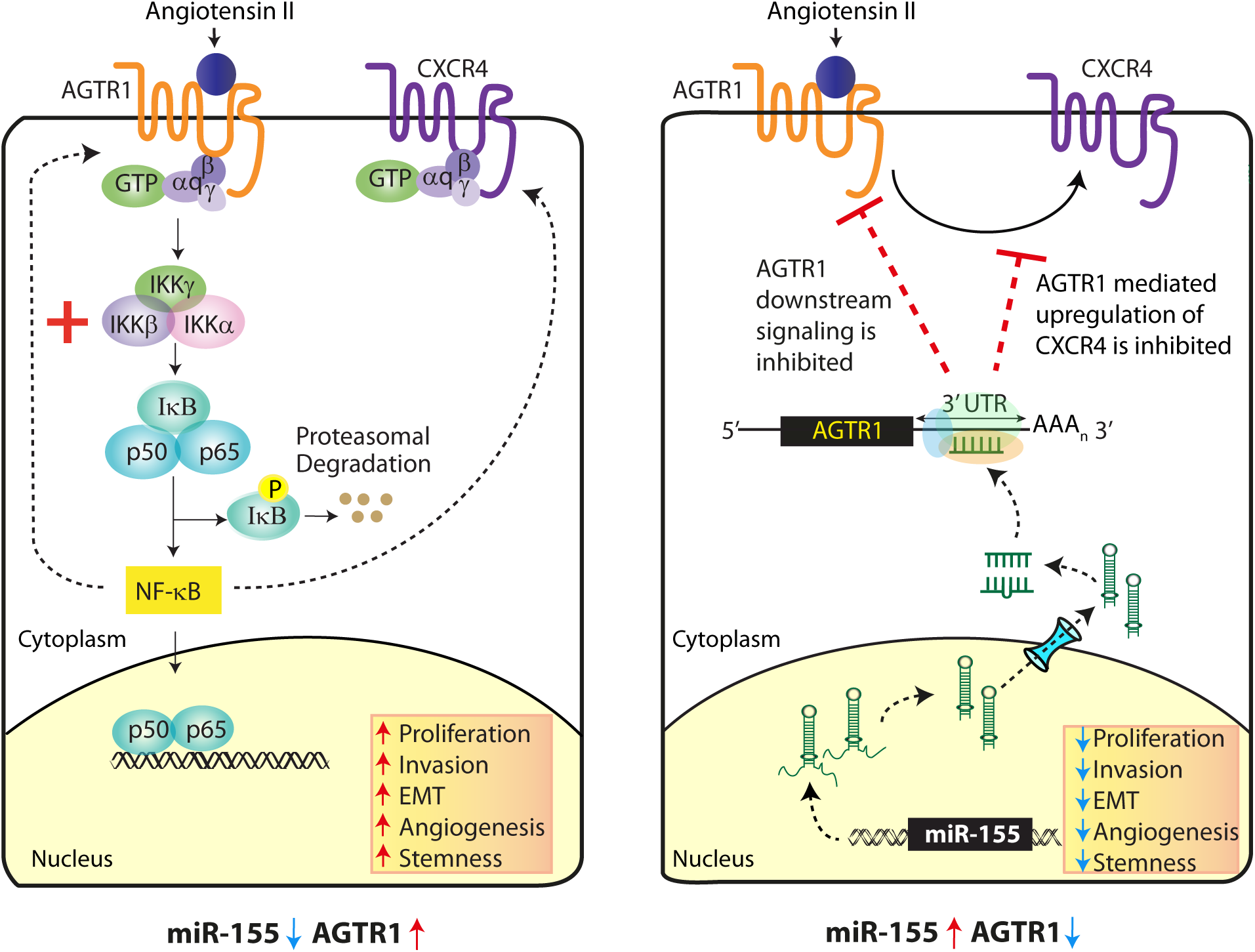
MiR-155 mediated AGTR1 downregulation leads to reduced NF-κB signaling and CXCR4 levels attenuating their associated oncogenic properties. Schematic representation of angiotensin II stimulated, AGTR1 downstream signaling leading to activation of NF-κB pathway which in turn increases CXCR4 expression and mediates feedback upregulation of AGTR1 (left panel). While miR-155 post-transcriptionally downregulates AGTR1 which leads to reduced NF-κB signaling, decrease in CXCR4 expression and inhibition of feedback regulatory loop. Collectively, miR-155 disrupts the oncogenic NF-κB/CXCR4/AGTR1 signaling axis in GBM thereby decreasing AGTR1-mediated tumorigenesis.

## Discussion

Elevated levels of AGTR1 have been associated with high grade malignancy, angiogenesis, and poor survival in glioma [7, 12]. However, the underlying molecular mechanism involved in AGTR1 upregulation in GBM still needs to be understood. Here, we have uncovered the regulatory mechanism involved in AGTR1 overexpression. We demonstrated that miR-155 post-transcriptionally regulates *AGTR1* expression in GBM and elicits pleiotropic anticancer effects such as inhibition of angiogenesis in CAM assay and significant regression of tumor burden in xenograft mice model. Although, miR-155 has been implicated as a probable oncogenic miRNA in multiple cancers [42, 43], its tumor-suppressive role has also been observed, wherein it inhibits the invasive as well as metastatic potential in colorectal carcinoma [44]. In BCa, miR-155 directly attenuates the expression of TCF4 transcription factor, an important regulator of EMT [45]. Moreover, miR-155 is also known to inhibit HER2 expression by post-transcriptional regulation as well as by targeting Histone Deacetylase 2 (HDAC2), its transcriptional activator [46]. In triple negative breast cancer, high miR-155 levels are known to be associated with low recombinase RAD51 levels and increased patient survival [47]. Moreover, miR-155 is also known to reduce the CD44+/CD117+ ovarian cancer-initiating cells which display stem cell-like properties by targeting *CLDN1* [48] and inhibit cancer stem cell like properties in colorectal cancer by targeting TGFβ/SMAD signaling pathway [49]. In conjunction with these studies, we observed that miR-155 inhibits the oncogenic potential of GBM by inhibiting EMT, stemness and promoting cell death.

In human umbilical vein endothelial cells, miR-155 is shown to target *AGTR1*, leading to decreased ATII induced ERK1/2 phosphorylation [50]. Similarly, we also observed a decrease in ERK1/2, MAPK signaling, mesenchymal markers along with an increase in apoptosis in miR-155 overexpressing GBM cells. Earlier, it has been known that miR-155 exerts anti-angiogenic effects by suppression of AGTR1 and Suppressor Of Cytokine Signaling 1 (SOCS-1) [51]. These studies corroborate with our findings wherein miR-155 overexpressing GBM cells fail to induce prominent angiogenesis compared to control in chick CAM angiogenesis assay. Of late, it has been reported that miR-155 mediated gene regulation is dependent on the cellular context [52], and the phenotype associated with miR-155 targeting the same transcript in different cell types will probably be different [53]. However, the results from our study establish the tumor-suppressive role of miR-155 in AGTR1-positive subtype of GBM, wherein, its overexpression results in inhibition of multiple oncogenic properties and signaling pathways.

The NF-κB plays a major role in cancer development by dysregulation of IKK activity leading to cell proliferation, angiogenesis, migration, and metastasis in multiple cancers including GBM [54–56]. Moreover, AGTR1 is known to mediate its oncogenic effects in BCa through activation of NF-κB signaling [17]. It has also been established that NF-κB subunits p65 and p50 directly binds to the *CXCR4* promoter, and activate its expression, leading to increased cell migration and metastasis [21]. Interestingly, ATII stimulated murine neuronal cells also exhibit a positive feedback mechanism of AGTR1 upregulation via NF-κB and CREB activation [40]. Furthermore, inhibiting NF-κB activity by using a NEMO (NF-κB essential modifier)–binding domain (NBD) peptide, depletion of IκB kinase 2 (IKK2), expression of a IκBαM super repressor, or by targeting inducible NF-κB genes is considered as an attractive therapeutic approach in GBM [57]. Similarly, it has been shown that overexpression of IκB, an inhibitor of κB in triple negative breast cancer cells, led to reduced expression of CXCR4 [21], and mutant IκBα super-repressor mediated repression of NF-κB in prostate cancer results in the decrease of CXCR4 expression, adhesion and transendothelial migration [58].

Recently, it has been shown that AGTR1 overexpression in BCa engages both ligand-independent and -dependent activation of NF-κB signaling through a triad of CARMA3, Bcl10, and MALT1, known as CBM signalosome [17]. Moreover, they also demonstrated that CBM-dependent activation of NF-κB promotes tumor angiogenesis by inducing cancer cell-extrinsic effects. In the current study, we found that ATII stimulation results in the activation of NF-κB signaling pathway, subsequently resulting in increased CXCR4 and AGTR1 levels. Similarly, overexpression of miR-155 in GBM cells results in the downregulation of AGTR1 via post-transcriptional regulation, which in turn attenuates the NF-κB signaling pathway thereby leading to reduced CXCR4 and AGTR1 levels. Alternatively, we also demonstrated that pharmacological inhibition of NF-κB signaling pathway by an IKK inhibitor attenuates the expression of CXCR4 and AGTR1 in GBM. Moreover, SNB19 cells with higher levels of miR-155 show enhanced sensitivity to IKK-16 compared to control cells. Taken together, our study highlights that disrupting the AGTR1/NF-κB/CXCR4 axis via miR-155 and/or IKK inhibitor would be an effective therapeutic approach for the AGTR1-positive GBM patients.

In view of the pleiotropic anticancer effects displayed by miR-155 along with attenuation of the oncogenic AGTR1/NF-κB/CXCR4 axis, we propose that miRNA-155 replacement therapy could be explored for AGTR1-positive subtype of GBM. Nevertheless, the stability and accurate *in vivo* delivery of miRNAs still present challenges to translate it to the clinics [59]. Although, magnetic resonance guided focused ultrasound has been used to deliver miRNAs across blood brain barrier as a treatment modality for GBM [60] hence well-planned clinical trials with miR-155 are highly warranted for GBM patient with higher AGTR1 levels. Moreover, several strategies including small molecule inhibitors against IKK and its related kinases have been tested in preclinical studies [68, 71] as well as clinical trials for other diseases (Identifier: NCT00883584) and might be introduced for GBM after complete evaluation in clinical trials. Therefore, we also propose the use of IKK inhibitors to abrogate NF-κB signaling and disrupt the AGTR1/NF-κB/CXCR4 axis.

## Materials and Methods

### Xenograft model

NOD/SCID (NOD.CB17-Prkdcscid/J) mice of about five to six weeks old were randomly placed in two groups. The mice were subjected to anaesthesia with a cocktail of xylazine/ketamine (5 and 50 mg/kg, respectively) through the intraperitoneal route. Thereafter, SNB19-CTL and SNB19-miR-155 cells (5×10^6^ cells for each condition), resuspended in 100µl saline and mixed with 20% Matrigel were injected into dorsal flank of mice on both the sides. Digital Vernier’s calipers were used to measure tumor growth twice a week in a blinded assessment, and the formula (π/6) (L×W^2^), (L= length; W= width) was used to calculate the tumor volume. All procedures involving animals were approved by the Committee for the Purpose of Control and Supervision of Experiments on Animals (CPCSEA) and were in accordance with the guidelines of the Institutional Animal Ethics Committee at Indian Institute of Technology Kanpur.

### Luciferase Promoter reporter assay

pEZX-MT01 Dual-Luciferase reporter vector (*Renilla/Firefly*) from GeneCopoeia was used to clone a 525 bp region of *AGTR1* 3’UTR from human genomic DNA. Another similar region with mutated residues in the binding site of miR-155 was also cloned in the luciferase vector. SNB19 cells at a confluency of 30-40% were co-transfected with 25 ng pEZX-MT01 wild type and mutant constructs and 30pmol of miR-155 mimics using Lipofectamine RNAiMax (Invitrogen) for two consecutive days. Thereafter, the luciferase assay was terminated using the Dual-Glo Luciferase assay kit (Promega) following the manufacturer’s instructions. Normalization of Firefly Luciferase activity to Renilla luciferase activity was carried out for every sample analyzed [61].

### Gene expression array analysis

For gene expression profiling studies, RNA extracted from stable SNB19-CTL and SNB19-miR-155 cells was subjected to Agilent Whole Human Genome Oligo Microarray profiling (dual color) using Agilent Platform (8×60k format) in accordance with the manufacturer’s protocol. Two separate microarray hybridizations were performed using SNB19-miR-155 cells against the control SNB19-CTL cells. Locally weighted linear regression (Lowess) normalization was used to normalize the microarray data. To recognize significant gene expression patterns for differentially regulated genes, Pearson correlation coefficient-based hierarchical clustering algorithm was utilized. To identify differentially expressed genes, Benjamini and Hochberg procedure was used to calculate FDR- corrected *P*-values (FDR< 0.05). Genes with differential expression (log_2_ (fold change) > 0.6 or < -0.6, *P*< 0.05) were further analyzed using Database for Annotation, Visualization and Integrated Discovery (DAVID) for major deregulated pathways [62]. Gene set enrichment analysis (GSEA) was used for the identification of enriched molecular signatures in control with respect to miRNA overexpression [63]. Enrichment Map for critical biological pathways and processes was generated using Enrichment Map, a plug-in for Cytoscape network visualization software (http://baderlab.org/Software/EnrichmentMap/). Heatmap.2 function of ‘gplots’ in R was used to generate heatmaps.

### Western Blot Analysis

Protein samples were analyzed by estimating the cell lysates using the BCA protein estimation kit (G-Biosciences) and subjecting them to SDS-PAGE followed by transfer onto polyvinylidine difluoride (PVDF) membrane (GE Healthcare). Blocking was done in 2% bovine serum albumin (BSA) or 5% non-fat dry milk (NFDM) prepared in 1X tris-buffered saline with 0.1% Tween 20 (TBS-T) for an hour at room temperature (RT). The membrane was incubated overnight at 4°C with the following primary antibodies : 1:1000 diluted anti-AGTR1 (Abcam, ab124734), 1:1000 diluted p-Akt (CST, 2965), 1:1000 diluted total Akt (CST, 9272), 1:1000 diluted p-ERK (CST, 4377), 1:1000 diluted ERK (CST, 4695), 1:1000 diluted Bcl-xl (CST, 2764), 1:1000 diluted Bcl2 (CST, 762870), 1:1000 diluted E-cadherin (CST, 3195), 1:1000 diluted N-cadherin (CST, 4061), 1:1000 diluted Vimentin (CST, 3932), 1:5000 diluted β-actin (Abcam, ab6276), 1:1000 diluted phospho-NF-κB p65 (CST, 3033), 1:1000 diluted NF-κB p65 (CST, 4764), 1:1000 diluted CXCR4 (ab181020). Consequently, the blots were washed with 1X TBS-T followed by incubation with secondary anti-rabbit or anti-mouse antibody conjugated with horseradish peroxidase (Jackson ImmunoResearch, USA) for each respective antibody in 2% NFDM or BSA at RT for 2 hours. Finally, the membrane was washed and subjected to Pierce™ ECL Western Blotting Substrate (ThermoFisher) to visualize signals as per the manufacturer’s protocol.

### Chick Chorioallantoic Membrane (CAM) Angiogenesis assay

Fertilized eggs were incubated at 37°C with 70% humidity. A small hole was made on the blunt side of the egg and approximately 3ml of albumin was aspirated out from each egg with a needle followed by forming a small window on top of each egg which was later secured by scotch tape. The eggs were returned to the incubator until day six [64] when 3X10^6^ cells (SNB19-CTL and SNB19-miR-155) were implanted onto the CAM. The cells were resuspended in saline and mixed with 20% Matrigel. Subsequently, the windows were sealed properly before returning the eggs to the incubator. Three days post implantation, the eggs were taken out and the seal was removed. The cells were visualised in stereomicroscope under fluorescent light to detect their location as the cells have red fluorescence and then the image was taken in bright field via the microscope. Following this, the CAM was moistened with saline and fixed with 4% paraformaldehyde (PFA). The fixed CAM was then resected out of the egg and imaged under the stereomicroscope at 1.8X. The images obtained from both control (n=8) and experimental (n=8) set of eggs were analyzed by the Angiogenesis analyzer for ImageJ [37].

### Statistics

Two-tailed Student’s *t*-test was employed to assess the statistical significance of two distinct samples or indicated otherwise. The experimental groups were ascertained to be significantly different based on the *P*-value. Significance was determined as follows: \**P≤* 0.05, \*\**P≤* 0.001. Error bars depict the standard error of mean (SEM) acquired from three independent experiments.

## Supplementary methods and tables

Further methodologic details can be found in Supplementary Methods. Deregulated pathways on miR-155 expression can be found in Supplementary Table 1 and primers in Supplementary Table 2.

## Data availability

The gene expression microarray data from this study has been submitted to the NCBI Gene Expression Omnibus (GEO, http://www.ncbi.nih.gov/geo/) under the accession number GSE138737.

## Acknowledgments

Research funding from the Wellcome Trust/ DBT India Alliance (IA/I(S)/12/2/500635), Department of Biotechnology (BT/PR8675/GET/119/1/2015) and Science and Engineering Research Board, Government of India, (EMR/2016/005273) are acknowledged. AS thanks Indian Council of Medical Research, India for the Senior Research fellowship (ICMR number: 3/1/3/JRF-2011/HRD-88). We thank the Indian Institute of Technology Kanpur for the financial and infra-structure support. We also thank Dr. Jonaki Sen for extending the use of fertilized eggs’ facility and microscopy setup. Authors thank Vipul Bhatia, Ayush Praveen, Ritika Tiwari and Anjali Yadav for their technical assistance and insightful discussion. We also thank Vipul Bhatia, Ritika Tiwari, and Nishat Manzar for critically reading the manuscript.

## Authors’ Contributions

AS and BA designed and directed the experimental studies. AS performed the experiments and interpreted the results. AS and BA performed the mice xenograft studies. AS, NS, and BA interpreted and analyzed the data. AS and BA wrote the manuscript.

## References

1. Ostrom QT, Gittleman H, Fulop J, Liu M, Blanda R, Kromer C et al. CBTRUS statistical report: primary brain and central nervous system tumors diagnosed in the United States in 2008-2012. Neuro-oncology. 2015; 17: iv1-iv62.

2. Stupp R, Mason WP, Van Den Bent MJ, Weller M, Fisher B, Taphoorn MJ et al. Radiotherapy plus concomitant and adjuvant temozolomide for glioblastoma. New England Journal of Medicine. 2005; 352: 987–996.

3. Wick W, Osswald M, Wick A, Winkler F. Treatment of glioblastoma in adults. Therapeutic advances in neurological disorders. 2018; 11: 1756286418790452.

4. Behnan J, Finocchiaro G, Hanna G. The landscape of the mesenchymal signature in brain tumours. Brain. 2019; 142: 847–866.

5. Kim H, Verhaak RG. Glioma through the looking GLASS: molecular evolution of diffuse gliomas and the Glioma Longitudinal Analysis Consortium. 2018.

6. Ghosh D, Nandi S, Bhattacharjee S. Combination therapy to checkmate glioblastoma: clinical challenges and advances. Clinical and translational medicine. 2018; 7: 33.

7. Arrieta O, Pineda-Olvera B, Guevara-Salazar P, Hernández-Pedro N, Morales-Espinosa D, Cerón-Lizarraga T et al. Expression of AT1 and AT2 angiotensin receptors in astrocytomas is associated with poor prognosis. British journal of cancer. 2008; 99: 160.

8. Fogarty DJ, Sánchez-Gómez MV, Matute C. Multiple angiotensin receptor subtypes in normal and tumor astrocytes in vitro. Glia. 2002; 39: 304–313.

9. Fujimoto Y, Sasaki T, Tsuchida A, Chayama K. Angiotensin II type 1 receptor expression in human pancreatic cancer and growth inhibition by angiotensin II type 1 receptor antagonist. FEBS Lett. 2001; 495: 197–200.

10. Miyajima A, Kosaka T, Asano T, Asano T, Seta K, Kawai T et al. Angiotensin II type I antagonist prevents pulmonary metastasis of murine renal cancer by inhibiting tumor angiogenesis. Cancer Res. 2002; 62: 4176–4179.

11. Suganuma T, Ino K, Shibata K, Kajiyama H, Nagasaka T, Mizutani S et al. Functional expression of the angiotensin II type 1 receptor in human ovarian carcinoma cells and its blockade therapy resulting in suppression of tumor invasion, angiogenesis, and peritoneal dissemination. Clin Cancer Res. 2005; 11: 2686–2694.

12. Arrieta O, Guevara P, Escobar E, Garcia-Navarrete R, Pineda B, Sotelo J. Blockage of angiotensin II type I receptor decreases the synthesis of growth factors and induces apoptosis in C6 cultured cells and C6 rat glioma. Br J Cancer. 2005; 92: 1247–1252.

13. Rhodes DR, Ateeq B, Cao Q, Tomlins SA, Mehra R, Laxman B et al. AGTR1 overexpression defines a subset of breast cancer and confers sensitivity to losartan, an AGTR1 antagonist. Proceedings of the National Academy of Sciences. 2009; 106: 10284–10289.

14. Zhang Q, Yu S, Lam MMT, Poon TCW, Sun L, Jiao Y et al. Angiotensin II promotes ovarian cancer spheroid formation and metastasis by upregulation of lipid desaturation and suppression of endoplasmic reticulum stress. Journal of Experimental & Clinical Cancer Research. 2019; 38: 116.

15. Egami K, Murohara T, Shimada T, Sasaki K-i, Shintani S, Sugaya T, et al. Role of host angiotensin II type 1 receptor in tumor angiogenesis and growth. The Journal of clinical investigation. 2003; 112: 67–75.

16. Singh A, Srivastava N, Amit S, Prasad S, Misra M, Ateeq B. Association of AGTR1 (A1166C) and ACE (I/D) polymorphisms with breast cancer risk in North Indian population. Translational oncology. 2018; 11: 233–242.

17. Ekambaram P, Lee J-YL, Hubel NE, Hu D, Yerneni S, Campbell PG et al. The CARMA3–Bcl10–MALT1 Signalosome Drives NFκB Activation and Promotes Aggressiveness in Angiotensin II Receptor–Positive Breast Cancer. Cancer research. 2018; 78: 1225–1240.

18. McAllister-Lucas LM, Jin X, Gu S, Siu K, McDonnell S, Ruland J et al. The CARMA3-Bcl10-MALT1 signalosome promotes angiotensin II-dependent vascular inflammation and atherogenesis. Journal of Biological Chemistry. 2010; 285: 25880–25884.

19. Kim HJ, Hawke N, Baldwin AS. NF-κB and IKK as therapeutic targets in cancer. Cell death and differentiation. 2006; 13: 738.

20. Soubannier V, Stifani S. NF-κB signalling in glioblastoma. Biomedicines. 2017; 5: 29.

21. Helbig G, Christopherson KW, Bhat-Nakshatri P, Kumar S, Kishimoto H, Miller KD et al. NF-κ B promotes breast cancer cell migration and metastasis by inducing the expression of the chemokine receptor CXCR4. Journal of biological chemistry. 2003; 278: 21631–21638.

22. Schimanski CC, Schwald S, Simiantonaki N, Jayasinghe C, Gönner U, Wilsberg V et al. Effect of chemokine receptors CXCR4 and CCR7 on the metastatic behavior of human colorectal cancer. Clinical Cancer Research. 2005; 11: 1743–1750.

23. Zhou Y, Larsen PH, Hao C, Yong VW. CXCR4 is a major chemokine receptor on glioma cells and mediates their survival. Journal of Biological Chemistry. 2002; 277: 49481–49487.

24. Ehtesham M, Winston J, Kabos P, Thompson R. CXCR4 expression mediates glioma cell invasiveness. Oncogene. 2006; 25: 2801.

25. Ehtesham M, Mapara KY, Stevenson CB, Thompson RC. CXCR4 mediates the proliferation of glioblastoma progenitor cells. Cancer letters. 2009; 274: 305–312.

26. Stevenson CB, Ehtesham M, McMillan KM, Valadez JG, Edgeworth ML, Price RR et al. CXCR4 expression is elevated in glioblastoma multiforme and correlates with an increase in intensity and extent of peritumoral T2-weighted magnetic resonance imaging signal abnormalities. Neurosurgery. 2008; 63: 560–570.

27. Jiang Z, Zheng X, Rich KM. Down-regulation of Bcl-2 and Bcl-xL expression with bispecific antisense treatment in glioblastoma cell lines induce cell death. Journal of neurochemistry. 2003; 84: 273–281.

28. Rasper M, Schäfer A, Piontek G, Teufel J, Brockhoff G, Ringel F et al. Aldehyde dehydrogenase 1 positive glioblastoma cells show brain tumor stem cell capacity. Neuro-oncology 2010; 12: 1024–1033.

29. Pearson JR, Regad T. Targeting cellular pathways in glioblastoma multiforme. Signal transduction and targeted therapy 2017; 2: 17040.

30. Palanichamy K, Jacob JR, Litzenberg KT, Ray-Chaudhury A, Chakravarti A. Cells isolated from residual intracranial tumors after treatment express iPSC genes and possess neural lineage differentiation plasticity. EBioMedicine 2018; 36: 281–292.

31. Wick W, Brandes AA, Gorlia T, Bendszus M, Sahm F, Taal W et al. EORTC 26101 phase III trial exploring the combination of bevacizumab and lomustine in patients with first progression of a glioblastoma. American Society of Clinical Oncology, 2016.

32. Wen PY, Macdonald DR, Reardon DA, Cloughesy TF, Sorensen AG, Galanis E et al. Updated response assessment criteria for high-grade gliomas: response assessment in neuro-oncology working group. Journal of clinical oncology: official journal of the American Society of Clinical Oncology 2010; 28: 1963–1972.

33. Matsuda Y, Hagio M, Ishiwata T. Nestin: a novel angiogenesis marker and possible target for tumor angiogenesis. World journal of gastroenterology: WJG 2013; 19: 42.

34. Neves M, Fumagalli A, van den Bor J, Marin P, Smit MJ, Mayor F. The role of ACKR3 in breast, lung and brain cancer. Molecular pharmacology 2019: mol. 118.115279.

35. Roberts WG, Whalen PM, Soderstrom E, Moraski G, Lyssikatos JP, Wang H-F et al. Antiangiogenic and antitumor activity of a selective PDGFR tyrosine kinase inhibitor, CP-673,451. Cancer research 2005; 65: 957-966.

36. Hong X-Y, Wang J, Li Z. AGR2 expression is regulated by HIF-1 and contributes to growth and angiogenesis of glioblastoma. Cell biochemistry and biophysics 2013; 67: 1487–1495.

37. Carpentier G, Martinelli M, Courty J, Cascone I. Angiogenesis analyzer for ImageJ. 4th ImageJ User and Developer Conference proceedings, 2012, pp 198-201.

38. Yadav VN, Zamler D, Baker GJ, Kadiyala P, Erdreich-Epstein A, DeCarvalho AC et al. CXCR4 increases in-vivo glioma perivascular invasion, and reduces radiation induced apoptosis: A genetic knockdown study. Oncotarget 2016; 7: 83701.

39. Esencay M, Newcomb EW, Zagzag D. HGF upregulates CXCR4 expression in gliomas via NF-κB: implications for glioma cell migration. Journal of neuro-oncology. 2010; 99: 33–40.

40. Haack KK, Mitra AK, Zucker IH. NF-κB and CREB are required for angiotensin II type 1 receptor upregulation in neurons. Plos one. 2013; 8: e78695.

41. Waelchli R, Bollbuck B, Bruns C, Buhl T, Eder J, Feifel R et al. Design and preparation of 2-benzamido-pyrimidines as inhibitors of IKK. Bioorganic & medicinal chemistry letters. 2006; 16: 108–112.

42. Witten LW, Cheng CJ, Slack FJ. miR-155 drives oncogenesis by promoting and cooperating with mutations in the c-Kit oncogene. Oncogene. 2019; 38: 2151.

43. Zhou J, Wang W, Gao Z, Peng X, Chen X, Chen W et al. MicroRNA-155 promotes glioma cell proliferation via the regulation of MXI1. PloS one. 2013; 8: e83055.

44. Liu J, Chen Z, Xiang J, Gu X. MicroRNA-155 acts as a tumor suppressor in colorectal cancer by targeting CTHRC1 in vitro. Oncology letters. 2018; 15: 5561–5568.

45. Xiang X, Zhuang X, Ju S, Zhang S, Jiang H, Mu J et al. miR-155 promotes macroscopic tumor formation yet inhibits tumor dissemination from mammary fat pads to the lung by preventing EMT. Oncogene. 2011; 30: 3440.

46. He X, Zhu W, Yuan P, Jiang S, Li D, Zhang H et al. miR-155 downregulates ErbB2 and suppresses ErbB2-induced malignant transformation of breast epithelial cells. Oncogene. 2016; 35: 6015.

47. Gasparini P, Lovat F, Fassan M, Casadei L, Cascione L, Jacob NK et al. Protective role of miR-155 in breast cancer through RAD51 targeting impairs homologous recombination after irradiation. Proceedings of the National Academy of Sciences. 2014; 111: 4536–4541.

48. Qin W, Ren Q, Liu T, Huang Y, Wang J. MicroRNA-155 is a novel suppressor of ovarian cancer-initiating cells that targets CLDN1. FEBS letters. 2013; 587: 1434–1439.

49. Miao L. Mo1432–Inhibition of the Cancer Stem Cells-Like Properties by Mir-155, Involved in the Targeting of Transforming Growth Factor Beta/Smad2 Signal. Gastroenterology. 2019; 156: S-1306.

50. Cheng W, Liu T, Jiang F, Liu C, Zhao X, Gao Y et al. microRNA-155 regulates angiotensin II type 1 receptor expression in umbilical vein endothelial cells from severely pre-eclamptic pregnant women. International journal of molecular medicine. 2011; 27: 393–399.

51. Pankratz F, Bemtgen X, Zeiser R, Leonhardt F, Kreuzaler S, Hilgendorf I et al. MicroRNA-155 exerts cell-specific antiangiogenic but proarteriogenic effects during adaptive neovascularization. Circulation. 2015; 131: 1575–1589.

52. Hsin J-P, Lu Y, Loeb GB, Leslie CS, Rudensky AY. The effect of cellular context on miR-155-mediated gene regulation in four major immune cell types. Nature immunology. 2018; 19: 1137.

53. Michaille JJ, Awad H, Fortman EC, Efanov AA, Tili E. miR-155 expression in antitumor immunity: The higher the better? Genes, Chromosomes and Cancer. 2019; 58: 208–218.

54. Lee D-F, Hung M-C. Advances in targeting IKK and IKK-related kinases for cancer therapy. Clinical Cancer Research. 2008; 14: 5656–5662.

55. Xia Y, Shen S, Verma IM. NF-κB, an active player in human cancers. Cancer immunology research. 2014; 2: 823–830.

56. Robe PA, Bentires-Alj M, Bonif M, Rogister B, Deprez M, Haddada H et al. In vitro and in vivo activity of the nuclear factor-κB inhibitor sulfasalazine in human glioblastomas. Clinical Cancer Research. 2004; 10: 5595–5603.

57. Friedmann-Morvinski D, Narasimamurthy R, Xia Y, Myskiw C, Soda Y, Verma IM. Targeting NF-κB in glioblastoma: A therapeutic approach. Science Advances. 2016; 2: e1501292.

58. Kukreja P, Abdel-Mageed AB, Mondal D, Liu K, Agrawal KC. Up-regulation of CXCR4 expression in PC-3 cells by stromal-derived factor-1α (CXCL12) increases endothelial adhesion and transendothelial migration: role of MEK/ERK signaling pathway–dependent NF-κB activation. Cancer research. 2005; 65: 9891–9898.

59. Adams BD, Parsons C, Walker L, Zhang WC, Slack FJ. Targeting noncoding RNAs in disease. The Journal of clinical investigation. 2017; 127: 761–771.

60. Vega RA, Zhang Y, Curley C, Price RL, Abounader R. 370 magnetic resonance-guided focused ultrasound delivery of polymeric brain-penetrating nanoparticle microRNA conjugates in glioblastoma. Neurosurgery. 2016; 63: 210–210.

61. Bhatia V, Yadav A, Tiwari R, Nigam S, Goel S, Carskadon S et al. Epigenetic silencing of miRNA-338-5p and miRNA-421 drives SPINK1-positive prostate cancer. Clinical Cancer Research. 2019; 25: 2755–2768.

62. Huang DW, Sherman BT, Lempicki RA. Systematic and integrative analysis of large gene lists using DAVID bioinformatics resources. Nature protocols. 2008; 4: 44.

63. Subramanian A, Tamayo P, Mootha VK, Mukherjee S, Ebert BL, Gillette MA et al. Gene set enrichment analysis: a knowledge-based approach for interpreting genome-wide expression profiles. Proceedings of the National Academy of Sciences. 2005; 102: 15545–15550.

64. Deryugina EI, Quigley JP. Chick embryo chorioallantoic membrane models to quantify angiogenesis induced by inflammatory and tumor cells or purified effector molecules. Methods in enzymology. 2008; 444: 21–41.

