## Supplementary Figures and Methods for "Targeting AGTR1/NF-κB /CXCR4 axis by miR-155 attenuates oncogenesis in Glioblastoma"

**Financial support:** This work is supported by the extra-mural research grant from the Department of Biotechnology, Ministry of Science and Technology, Government of India [BT/PR8675/GET/119/1/2015 to BA].

**Conflict of interest:** The authors declare no conflicts of interest or disclosures.

***Corresponding Author**:
Bushra Ateeq, Ph.D.
Molecular Oncology Laboratory,
Department of Biological Sciences and Bioengineering,
Indian Institute of Technology Kanpur,
Kanpur, 208016, U.P., INDIA


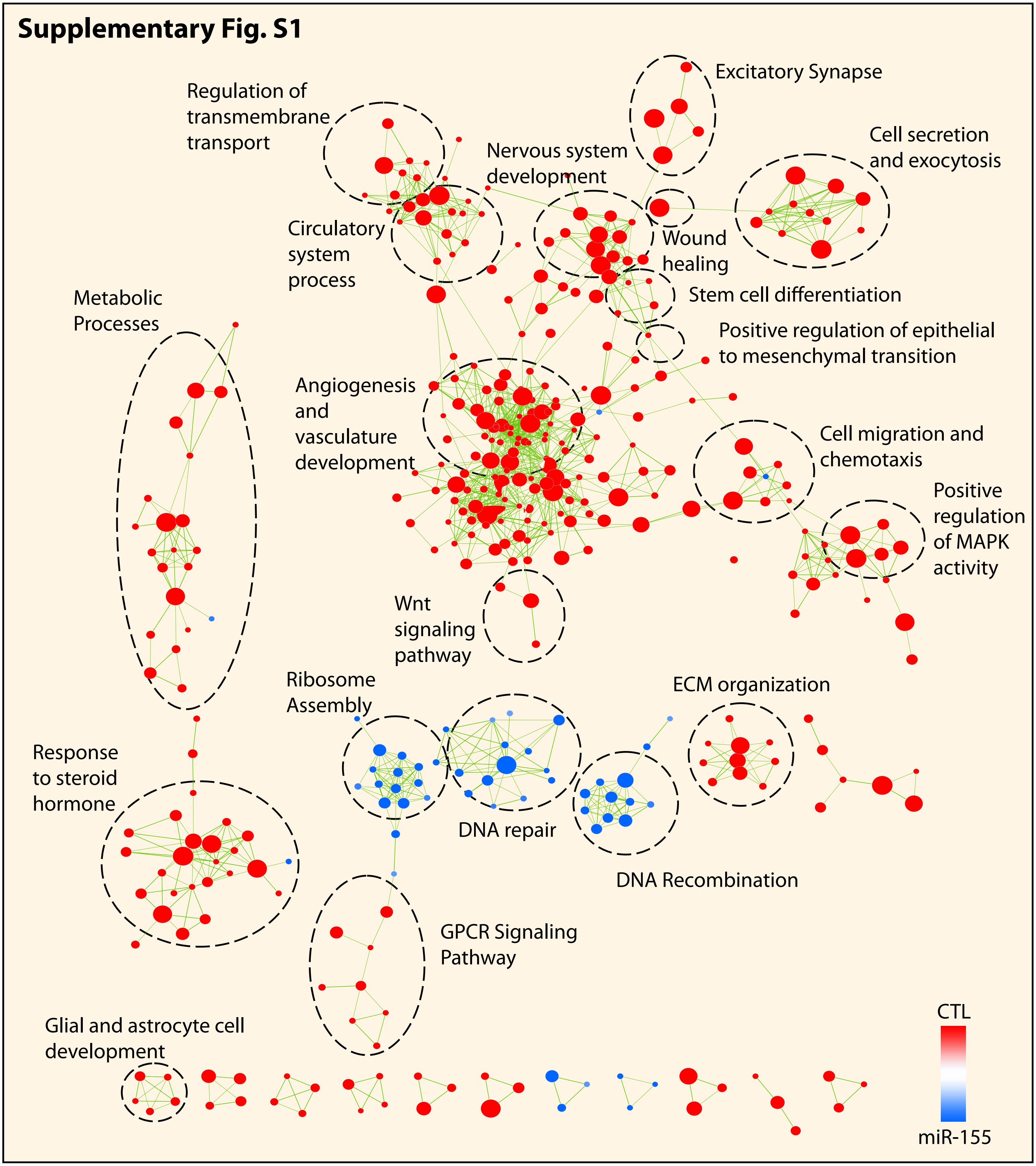


**Supplementary Fig. S1. Enrichment Map showing enriched biological pathways and processes.** Enrichment Map illustrating enrichment of pathways and processes including angiogenesis, cell migration, GPCR signaling and positive regulation of MAPK activity. Each node represents specific Gene Ontology (GO) term and node size indicates its significance. The connections between overlapping gene sets are denoted by green lines. Red color represents the pathways enriched in SNB19-CTL cells whereas blue represents pathways enriched in mir-155 overexpressing SNB19 cells. Clusters of genes that are functionally related to one another have been outlined by a dotted line and assigned a common label.

### Supplementary Fig. S2

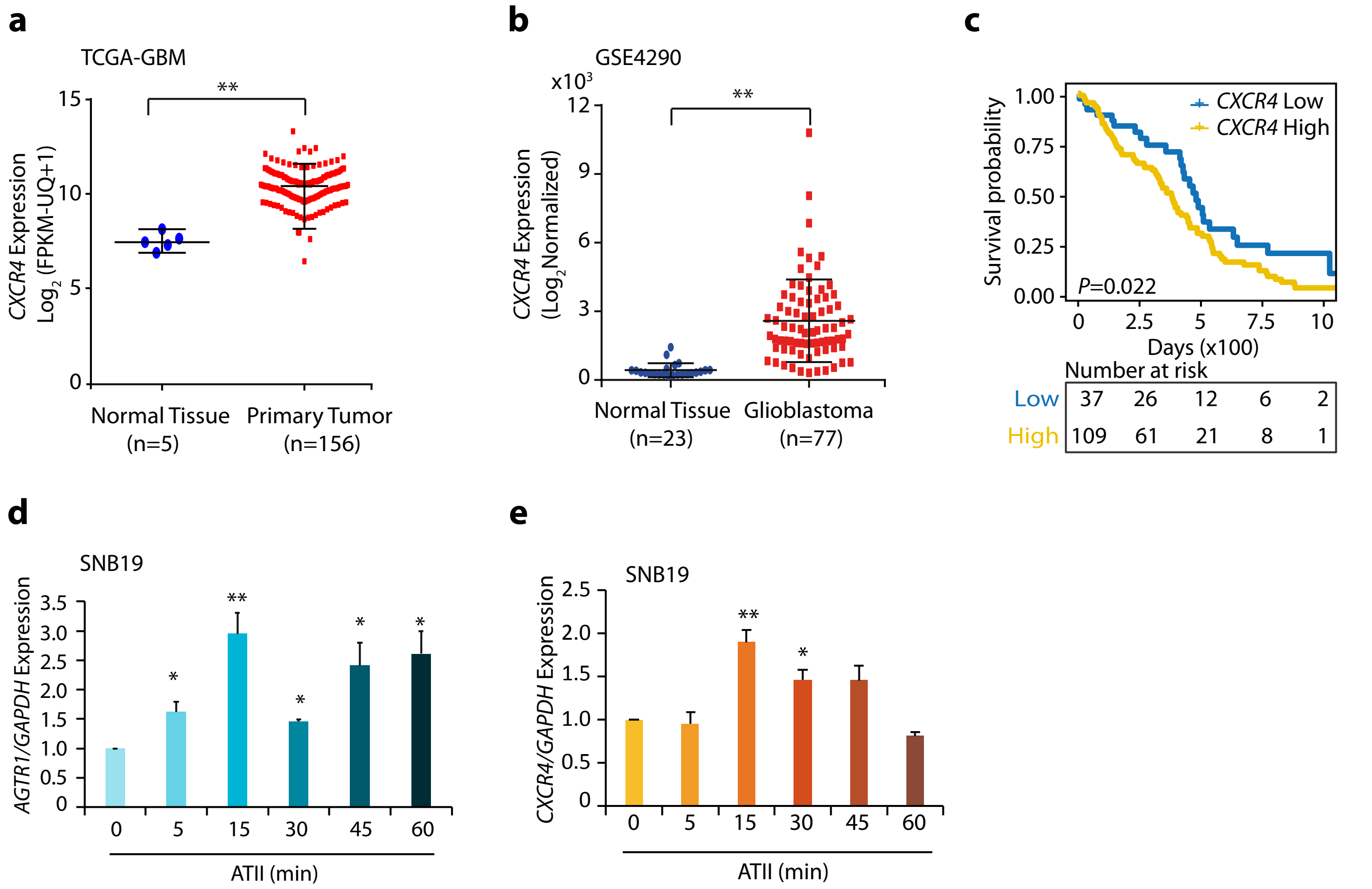


**Supplementary Fig. S2. CXCR4 is overexpressed in GBM patients and angiotensin II stimulation leads to increased AGTR1 and CXCR4 expression in GBM.**
**(a)** Scatter plot showing expression of *CXCR4* in normal tissue (n=5) and primary tumor (n=156) in TCGA-GBM cohort. **(b)** Scatter plot showing expression of *CXCR4* in normal tissue (n=23) and glioblastoma patients (n=77) in the Sun Brain cohort (GSE4290). **(c)** Kaplan-Meier curve showing survival probability of GBM patients with high versus low *CXCR4* levels in TCGA-GBM cohort. **(d)** Bar plot showing expression of *AGTR1* by quantitative PCR in SNB19 cells stimulated with Angiotensin II (ATII) for the indicated time points. **(e)** Same as **(d)** except, *CXCR4* expression was quantified. Data represent mean ± SEM. * $P \leq 0.05$, ** $P \leq 0.001$ using two-tailed unpaired Student's *t*-test.

**Supplementary materials and methods:**

**Cell lines and authentication**

The glioblastoma (SNB19 and U138) and human embryonic kidney 293 (HEK293T and HEK293FT) cell lines were cultured in media recommended by American Type Cell Culture (ATCC), supplemented with 10% fetal bovine serum (FBS) (GIBCO) along with 0.5% Penicillin Streptomycin (Pen Strep) (Thermo Fisher Scientific); cultured in $CO_2$ incubator (Thermo Fisher Scientific) at 37ºC with 5% $CO_2$. The glioblastoma cell lines used in the study were a kind gift from Prof. K. Somsundaram and Dr. V. Radha, whereas HEK293T and HEK293FT cells were procured from ATCC. For authenticating the cell lines, short tandem repeat (STR) profiling was performed at DNA Forensics, New Delhi and University of Arizona Genetics Core. Routine checks of Mycoplasma contamination were performed using the PlasmoTest mycoplasma detection kit (InvivoGen).

**Establishing stable miRNA overexpressing cell lines**

Human pre-miR-155 (miRBase accession ID MI0000681) was amplified by Polymerase Chain Reaction (PCR) from human genomic DNA ligated to digested, pLemiR vector, lentiviral miRNA expression vector with turbo RFP (Open Biosystems). Sanger sequencing was employed to confirm positive sequences of pLemiR constructs with integrated miRNA. For establishing stable miR-155 overexpressing cells, SNB19 cells were transfected with 2µg of miRNA cloned pLemiR construct employing FuGENE HD Transfection Reagent (Promega) in accordance with the manufacturer's instructions. After 48 hours, the cells were subjected to 0.5µg/ml of puromycin and the selection was continued for 3 to 4 weeks. Subsequently, Taqman assay (described elsewhere) was employed to validate the pooled population of cells obtained after selection for overexpression of miR-155. From the miR-155 overexpressing population, single clones were obtained by seeding the pooled cells in 96 well plate and subjecting them to puromycin selection for around 3-4 weeks. Finally, the single clones obtained were screened for miR-155 overexpression using Taqman probes and *AGTR1* expression employing quantitative PCR (qPCR).

**Transfection of microRNA mimetics**

Synthetic miRCURY LNA (Lock Nucleic Acid) microRNA mimics for human miR-155 (Cat. No. 472490-001) and non-targeting control mimic (Cat. No. 479903-001) were employed in miRNA overexpression. Briefly, the cells seeded at 40% confluency were transfected the next day with miR-155 mimic. After 24 hours a second transfection was performed. The transfection was performed using Lipofectamine RNAiMAX (Invitrogen) with a final concentration of 30pmol. Further, 48 hours post transfection the cells were processed for the assay.

**Real Time Quantitative PCR**

MiRNeasy Mini Kit (Qiagen) was used for extracting total RNA for miRNA related experiments or else TRIzol (Ambion) was employed. 1μg of RNA with good integrity was subjected to cDNA synthesis using SuperScript III First Strand synthesis System (Invitrogen) in accordance with the manufacturer's instructions. All reactions were performed in triplicates for quantitative PCR (qPCR) using SYBR Green Master Mix (Applied Biosystems). For calculating the relative expression of the target gene, ΔΔCt method as previously described [1] was used with the primers listed in the Supplementary Table S2.To carry out qPCR for miRNA stem loop with target specific stem loop reverse transcription primers, TaqMan microRNA reverse transcription kit (Thermo Fisher Scientific) was used. Subsequently, Taqman assays (Applied Biosystems) were performed according to the manufacturer's instructions. Relative expression of target miR-155 (Applied Biosystems Assay ID: 000479) was normalized to RNUB6 (Assay ID: 4427975).

**Cell proliferation assay**

Cell proliferation assay for SNB19 CTL, Pool, C-1, C-2, and C-3 clones was performed by seeding $1x10^4$ cells per well of 12 well culture plates. Cells were trypsinized at specified time points and counted by the Z-series Coulter counter (Beckman Coulter). Alternatively, cell viability was measured by using a colorimetric assay for 96-well plates with cell proliferation reagent WST-1 (Roche), for accessing the effect of IKK-16 on cell viability. SNB19 cells ($2x10^3$) were treated with indicated concentrations of IKK-16 followed by measuring absorbance at 450nm post incubation in WST-1. For determining the effect of *AGTR1* knockdown on cell proliferation we used resazurin sodium salt solution (w/v; Sigma, St. Louis MO). SNB19 CTL and SNB19 shAGTR1(1 and 2) cells were plated at a confluency of $2x10^3$ cells per well of 96 well plate. 10µl of rezazurin was added to each well followed by incubating the plates for 4 hours in dark at 37ºC and measuring the fluorescence at (530/590 nm ex/em) by a plate reader.

**Migration and Invasion Assays**

For cell invasion assays, Transwell Boyden chambers of 8µm pore size (Corning) were used. Briefly, $1x10^5$ cells resuspended in serum free (SF) RPMI-1640 were added onto Matrigel (BD Biosciences) coated Transwell insert, whereas conditioned media obtained from the cells and supplemented with 10% FBS was added to the lower chamber to serve as a chemoattractant. The invasion was accessed after 24 hours incubation of cells at 37ºC along with 5% $CO_2$. The chambers were taken out, media was removed and left to dry overnight. Further, the chambers were stained with 0.5% (w/v) of crystal violet dissolved in methanol and diluted in 1X PBS. Representative fields were imaged in the Axio Observer ZI microscope (Zeiss). The quantification of invaded cells was done by de-staining with 10% (w/v) glacial acetic acid in distilled water, followed by measuring the absorbance of the de-staining solution at 550 nm. A similar protocol was employed for migration assay except no Matrigel was coated onto the chambers and the assay was terminated at 16 hours post incubation.

**Foci formation assay**

Foci formation assay was performed by plating 2x$10^3$ cells per well in a 6 well cell culture dish in RPMI-1640 supplemented with 5% FBS (Invitrogen) followed by incubation at 37ºC along with 5% $CO_2$, media was changed every two days. For IKK-16 foci assay, the media changed was supplemented with IKK-16 (2µM and 4µM) as indicated. The assay was terminated after two weeks and the foci were fixed in 4% paraformaldehyde (PFA) and stained in 0.05% (w/v) of crystal violet solution. The foci were either counted under the microscope manually field wise or the crystal violet was de-stained using 10% glacial acetic acid and absorbance measured at 550 nm.

**Soft agar colony formation assay**

To access anchorage independent growth, soft agar plates were prepared. Firstly, 2 ml of 0.6% low melting agarose (Sigma-Aldrich) in RPMI-1640 medium was poured into 6 well dishes and allowed to polymerize. Following this, a second layer of 2 ml 0.3% low-melting agarose in RPMI-1640 medium along with stable SNB19 CTL, Pool, C-1, C-2, and C-3 cells respectively, were poured onto the first layer. The assay plates were incubated at 37ºC along with 5% $CO_2$ for 20 days, and were supplemented with 2x media every third day after which the colonies were observed and imaged under a microscope.

**Lentiviral Packaging**

Virapower Lentiviral Packaging Mix (Invitrogen) was used to produce lentiviral particles as per the instructions provided by the manufacturer. Briefly, HEK293FT cells were plated in a 100mm cell culture dish at a confluency of 90% in media devoid of antibiotics. pLKO.1 shAGTR1 vectors along with the Virapower Lentiviral Packaging Mix were transfected using FuGENE HD Transfection reagent (Promega). The media was replaced 24 hours later and the viral particles were harvested after 48-60 hours by collecting the media, following which viral particles aliquots were made and stored in -80ºC. To generate stable cell lines, SNB19 cells were plated in a 6 well cell culture dish and infected with viral particles along with polybrene (hexadimethrine bromide; 8μg/ml) (Sigma-Aldrich). The media was replaced the next day and puromycin (Sigma-Aldrich, R0908) selection at a concentration of 1 μg/ml was started after three days of infection.

**Cell cycle analysis and Flow Cytometry**

For cell cycle analysis, SNB19 cells were transfected with miR-155 mimics or control RNA. Twenty four hours post transfection, cells were fixed with 70% ethanol and stained with Propidium Iodide (Cat. no. 421301, BioLegend) according to the manufacturer's instructions. Staining with CD44 antibody (Miltenyi Biotec, cat. no: 130-098-108, 1:50) was performed by incubating SNB19-miR155 and SNB19-CTL cells for one hour with the antibody at 4 ºC, followed by washing with 1X PBS twice and acquisition. The data was acquired on Beckman Coulter's CytoFLEX platform and analysed with FlowJo Software.

**ALDH assay**

Aldeflour assay was carried out to determine the enzymatic activity of aldehyde dehydrogenase (ALDH) using the Aldefluor kit (Stemcell Technologies) in accordance with the manufacturer's instructions. Briefly, the cells were washed with 1X PBS by centrifugation at 1500 rpm at 4ºC for

5 minutes. Following this, 1 ml of Aldefluor assay buffer was added to cells and the pellet was resuspended in it. Thereafter, 5 µl of activated Aldeflour substrate was added. The cell suspension was divided into two tubes. Where one tube was left as it is, 5µl of ALDH inhibitor, diethylaminobenzaldehyde (DEAB) was added to another tube. Further, the cells were incubated for 30 minutes at 37ºC followed by centrifugation and resuspension in 500 µl of Aldeflour assay buffer. The green channel was used to detect ALDH activity. For gating the ALDH+ population of cells, cells treated with DEAB were used as respective controls to their samples. Beckman Coulter's CytoFLEX platform was used for data acquisition and FlowJo Software was used for the analysis.

1 Tiwari R, Pandey S, Goel S, Bhatia V, Shukla S, Jing X *et al*. SPINK1 promotes colorectal cancer progression by downregulating Metallothioneins expression. *Oncogenesis* 2015; 4: e162.
